# Disentangling Production and Persistence of Extracellular Virions in Grassland Soils with SIP-Viromics

**DOI:** 10.1101/2025.05.25.655894

**Authors:** Gareth Trubl, Simon Roux, Matthew Kellom, Dariia Vyshenska, Andy Tomatsu, Kanwar Singh, Jeffrey A. Kimbrel, Emiley Eloe-Fadrosh, Rex R. Malmstrom, Jennifer Pett-Ridge, Steven J. Blazewicz

## Abstract

Viruses are abundant and ecologically important in soils, yet the persistence and production dynamics of extracellular virions remain poorly understood. We applied a genome-resolved stable isotope probing viromics (SIP-viromics) approach, combining H ^18^O labeling with viral metagenomics, to track virion turnover in seasonally dry grassland soils following rewetting. We identified 354 viral populations (vOTUs) using individual-sample and combined metagenome assemblies. Only 22% of vOTUs exhibited significant ^18^O enrichment, indicating active replication and new virion production during the 1-week incubation; the majority (78%) persisted without detectable replication, consistent with a viral seed bank. Active vOTUs accounted for 4.76–5.15% of total virions per gram of soil, with viral loads ranging from 3.15 x 10^10^ to 6.59 x 10^10^ virions per gram. Probabilistic and deterministic sensitivity analyses spanning viral DNA fraction and genome length reinforced that persistent virions represented the majority of the extracellular viral pool post-wet-up, regardless of parameter assumptions. Host predictions linked both active and persistent vOTUs primarily to Actinomycetota and Pseudomonadota—bacterial groups known to rapidly resuscitate following rewetting—suggesting that some viruses exhibit rapid turnover while others persist over longer timescales, forming a stable viral pool capable of reinitiating infections during favorable conditions. These results demonstrate that SIP-viromics can distinguish newly produced from persistent virions and reveal host-associated patterns of lytic infection and virion production. Our findings advance understanding of soil virus-host interactions and highlight the ecological role of persistent virions as a genetic reservoir contributing to microbial turnover and biogeochemical cycling following environmental disturbance.

**Importance:** Understanding the persistence and production dynamics of soil viruses is critical for elucidating their roles in microbial community dynamics and nutrient cycling, yet these processes have remained largely uncharacterized due to methodological limitations. By integrating stable isotope probing with viromics, this study provides a robust framework for directly distinguishing newly produced from persistent virions *in situ*. Unlike conventional viromics, which only catalogs viral diversity, SIP-viromics enables quantification of active viral replication and persistence under natural soil conditions. Our results demonstrate that most virions in a seasonally dry soil persisted through a rewetting event, with active replication limited to a minority of viral populations. Persistent virions were primarily linked to dominant bacterial groups, indicating that host ecophysiology and environmental stability strongly influence lytic infection. Collectively, these findings highlight viruses as long-term reservoirs of genetic material, capable of shaping microbial dynamics and ecosystem processes over time. This work establishes SIP-viromics as a powerful approach for studying virus-host interactions and their ecological significance in terrestrial environments.

## Introduction

Isotope probing is a powerful tool for microbial ecology, enabling *in situ* tracing of metabolic activity and ecological interactions by monitoring isotope incorporation into biomolecules. Historically, isotope tracing methods have been widely used in microbial and viral cultures to examine tracer uptake into biomolecules (e.g., DNA, RNA, and lipids) and investigate specific metabolic pathways (Tang et al., 2009; Radajewski et al., 2000; Abraham, Hesse, and Pelz, 1998). These approaches also helped establish foundational principles in biology, including the role of DNA as hereditary material (Hershey et al., 1951) and key mechanisms of viral reproduction and virus-host interactions (Putnam and Kozloff, 1948; Putnam and Kozloff, 1950; Kozloff et al., 1951; Putnam et al., 1952; Ngo et al., 2024). More recently, stable isotope probing (SIP) has been coupled with whole-community DNA sequencing (shotgun metagenomics) and a mathematical model to quantify isotope enrichment (e.g., quantitative SIP; Hungate et al., 2015) to study complex environments (Greenlon et al., 2022), expanding the application of isotope probing to enable precise, genome-resolved tracking of metabolic activity and ecological interactions within microbial communities.

This expanded application has enabled the *in situ* detection of actively replicating viruses and highlighted their roles in host turnover and carbon cycling. However, most SIP-enabled studies have primarily focused on intracellular viral DNA (Lee et al., 2021, 2023; Barnett and Buckley, 2023), leaving open questions about the fate and function of extracellular virions. In particular, the persistence of unlabeled, potentially ‘inactive’ virions remains poorly understood. Recent work has shown that viral biomass can increase substantially following rewetting of seasonally dried soils, contributing significantly to microbial mortality and CO₂ efflux (Nicolas et al., 2023). Yet, it is unclear which virions are newly produced versus those persisting in the environment. This distinction is important for understanding viral survival strategies, turnover, persistence during environmental perturbation, host-range dynamics, and their broader impacts on biogeochemical cycling.

Despite the known ecological importance of soil viruses, several key knowledge gaps remain unresolved. We currently do not know which fraction of the extracellular virion pool was recently produced versus what fraction represents a persistent, pre-existing reservoir (or viral seed bank) in general. In addition, the viability of this persistent pool—and how it depends on host life history traits and virion structural properties—remains unclear. As a result, we cannot yet resolve how survival strategies, turnover rates, and persistence mechanisms differ across host taxa in seasonally dry soils or how persistent virions shape post-disturbance community dynamics and nutrient cycling.

To address this gap, we applied SIP-viromics, a novel approach that integrates SIP with viromics to track extracellular virion dynamics in soil. Unlike whole-community metagenomics, which captures both intra- and extracellular viruses, SIP-viromics specifically targets the extracellular virus fraction by concentrating and purifying virions from environmental samples. Because oxygen from water is universally required for DNA synthesis, ^18^O-SIP enables the estimation of total viral production and persistence, rather than substrate-specific activity. By measuring ^18^O incorporation into virion DNA following wet-up in a seasonally dry California grassland, we were able to distinguish newly produced virions from those that persisted through dry conditions without replicating, providing insights into viral turnover, survival strategies, and ecological dynamics. Building on prior SIP advances (Nuccio et al., 2022; Vyshenska et al., 2023), this approach extends traditional viromics—which recovers greater viral diversity than whole-community metagenomics (Trubl et al., 2018; Ter Horst et al., 2021; Santos-Medellín et al., 2021)—by enabling the differentiation of newly replicated and persistent virions. This capacity to disentangle viral production and persistence is broadly transferable to other ecosystems, including aquatic environments and the human gut, where understanding viral replication dynamics and virion persistence is central to interpreting virus-host interactions and ecosystem impacts. The SIP-viromics workflow includes isotope incubation, viral particle isolation and purification, DNA extraction, density-based separation of labeled DNA, and high-throughput sequencing (Fig. 1).

**Figure 1.**
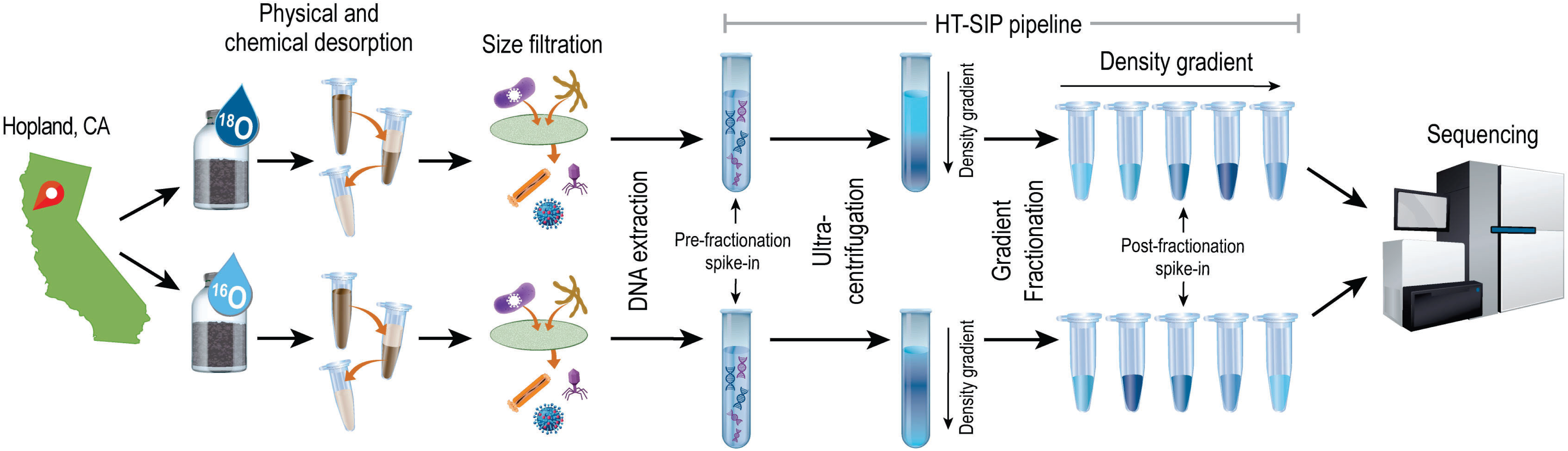
Overview of SIP-virome experimental workflow. Dry soil was collected from Hopland, California, and incubated in jars amended with either ^18^O-enriched water (H ^18^O; referred to as ^18^O) or natural abundance water (H ^16^O; referred to as ^16^O). After one week of incubation, a virome extraction was performed on the soils, where virions were physically and chemically desorbed from the soil matrix and isolated via size filtration. We then extracted DNA from the purified virions, separated it by density using ultracentrifugation, collected individual density fractions, and sequenced them.

## Results and Discussion

### Viral population recovery and prevalence

Using our SIP-viromics method, we identified 354 viral populations (vOTUs) through two types of metagenome assembly approaches: individual assembly of each sample and a combined assembly of all libraries (Table S1). Of these vOTUs, 165 were exclusive to the combined assembly, 91 were unique to individual sample assemblies, and 98 were shared between both approaches (Fig. S1). This highlights how single-sample and combined assemblies are both useful to assemble long (≥10 kb) virus genome fragments from this type of metagenome. To evaluate vOTU distribution across samples, we tested two horizontal genome coverage thresholds for detection. Using a relaxed 10% minimum horizontal coverage cutoff, 354 vOTUs were present and 82.2% of vOTUs were detected across all four samples. This value dropped to 50.5% when using a stricter 75% cutoff and only 274 vOTUs were detected (Fig. S1).

While higher thresholds reduce the risk of false positives, they can underestimate vOTU prevalence, particularly in viromes subjected to multiple rounds of amplification and characterized by uneven genome coverage. Previous work in viral ecology has shown that relaxed cutoffs are often more appropriate for detecting low-abundance viruses in such datasets (Paez-Espino et al., 2016; Trubl et al., 2019). Based on this context and the goal of capturing the breadth of the viral community, we applied the 10% cutoff for downstream vOTU analyses. Consistent with the virome-enrichment goal of our workflow, ViromeQC (Zolfo et al., 2019) analysis showed low rRNA marker alignment rates across libraries (SSU rRNA: 0.006–0.20%; LSU rRNA: 0.008–0.28%) and low bacterial marker alignment rates (0.045–1.32%), indicating that non-viral cellular DNA was a minor component of these viromes (Table S2).

Rewetting of seasonally dried soils triggered a sharp increase in microbially driven CO_2_ production (Fig. S2: Table S3), indicating heightened microbial activity and likely increases in host-virus interactions during this time. The moisture content used in this experiment (30% gravimetric water content) corresponds to approximately 60–70% water-filled pore space, a range widely recognized as supporting maximal microbial activity in non-saturated, aerobic soils (Linn and Doran, 1984; Skopp et al., 1990). Below this range, metabolic activity is increasingly constrained by limited pore connectivity and diffusional limitation of substrates, whereas moisture levels above this range reduce oxygen diffusion and can suppress aerobic microbial processes (Or et al., 2007; Manzoni et al., 2012). Thus, 30% gravimetric water content reflects hydrologically realistic post-first-rain conditions under which these xeric grassland soils typically experience rapid microbial resuscitation and elevated respiration (Placella et al., 2012; Barnard et al., 2013). Nevertheless, soil moisture exerts strong control over nutrient diffusion, host-virus encounter rates, and DNA replication, and wetter soils, or larger hydrologic pulses, may support higher viral activity than observed here. In more mesic systems, or under greater wet-up magnitude, a larger fraction of the viral community may become active as host growth increases and physical connectivity within the soil matrix improves. Conversely, in strongly water-limited environments, persistent virions may dominate for longer periods. Therefore, while our results demonstrate that persistent virions form the majority of the extracellular viral pool under hydrologically realistic conditions for this xeric system, we do not interpret this pattern as universal across soil types. Future SIP-viromics work across soils differing in texture, nutrient status, and hydrologic regimes will be essential for determining how moisture availability shapes viral activation, persistence, and their ecological consequences.

Approximately 78% of virions from vOTUs did not exhibit significant ^18^O enrichment, indicating that the majority of viral populations either did not replicate or did so at levels below detection during the week following rewetting (Fig. 2; Table S4). These virions were likely present in the dry soil prior to rewetting and remained stable throughout the incubation period. This limited activity is consistent with drought-constrained soil environments, where reduced pore connectivity and increased solute concentrations can restrict microbial processes and associated virus-host interactions (Shan et al., 2026). Such stability suggests that virion turnover in our microcosms occurred on weekly or longer timescales. If these virions remain infectious, their persistence could represent a survival strategy and help explain the consistently high extractable virus-like particle (VLP) counts observed in soils (10^7^–10^10^ VLPs per gram; Williamson et al., 2017; Cao et al., 2022). Conversely, the remaining 22% of viral populations showed significant ^18^O enrichment, providing direct evidence of replication and release of new virions during the rewetting period (Fig. 2; Fig. S3; Table S4). Successful ^18^O incorporation into viral DNA and its detection in purified virions validates the SIP-viromics approach. Across samples, active vOTUs generally did not differ in relative abundance from persistent ones, and in two cases, persistent vOTUs were more abundant (Wilcoxon test, p < 0.05; Table S5). This suggests that vOTUs active in bulk soil following wet up, represent a minority of the viral community and are preferentially associated with microbial resuscitation.

**Figure 2.**
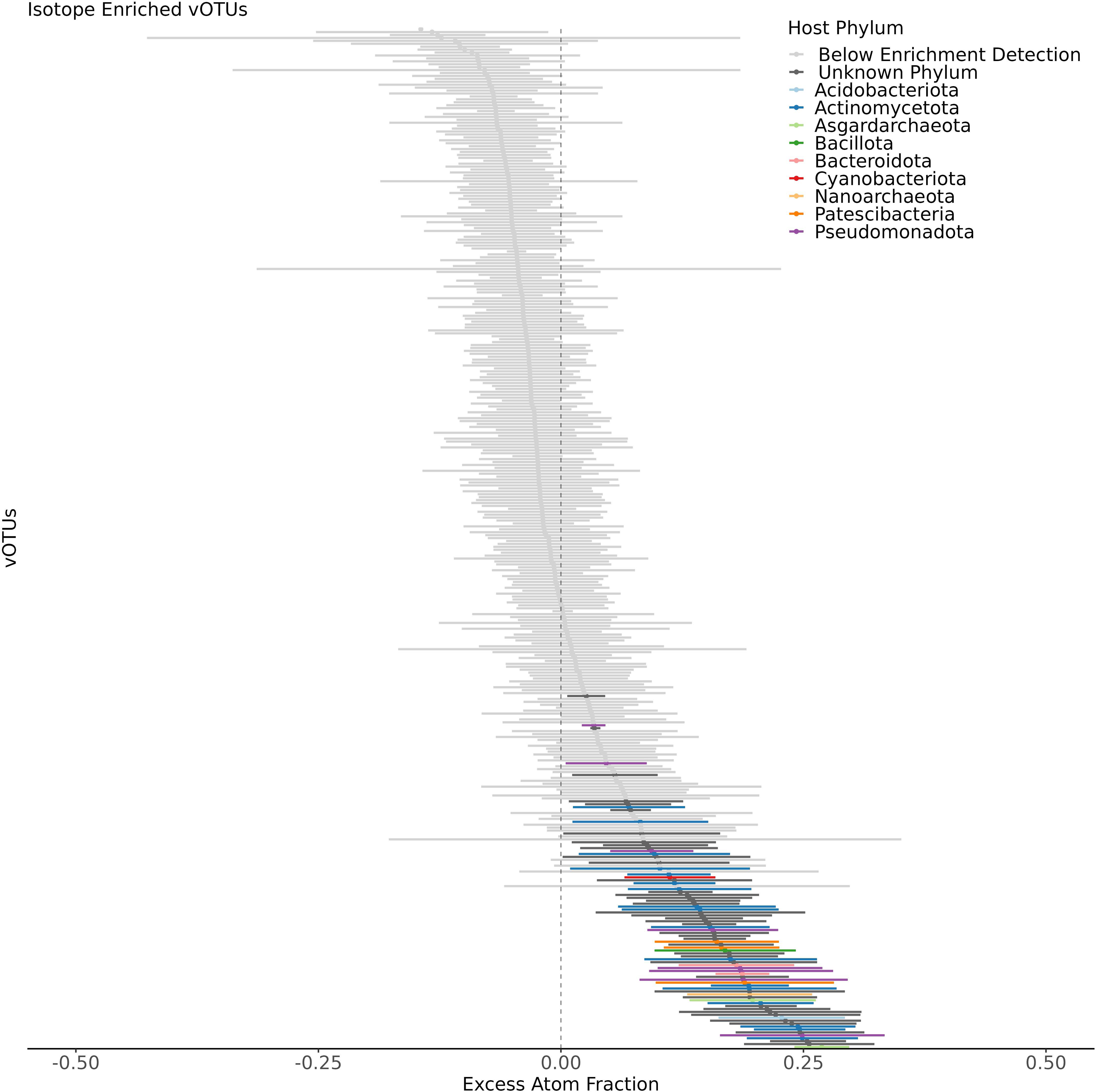
Estimated ¹⁸O enrichment values for viral populations and predicted host for active viral populations. Viral populations (vOTUs) are ordered from lowest to highest excess atom fraction (EAF) (top to bottom). Points represent EAF estimates based on density-resolved viral abundance across 16 DNA density fractions, using two biological replicates per treatment (H ^18^O-labeled and unlabeled controls). Horizontal lines indicate 95% confidence intervals (CI) derived from isotope incorporation model fits. vOTUs with detectable ^18^O enrichment (lower CI > 0) are shown with solid confidence interval lines and colored by their predicted host phylum (black for unknown hosts). vOTUs without detectable enrichment (lower CI < 0) are shown in gray. The relative abundance of active vOTUs was compared to persistent vOTUs using a Wilcoxon test, which showed persistent vOTUs were significantly more abundant in two of the biological samples (p < 0.05; see Table S5).

### Quantification of active and persistent virions

To quantify viral activity, we estimated virions per gram of soil across samples and mapping thresholds, distinguishing active, newly produced virions from persistent vOTUs. Total virion counts ranged from 3.15 x 10^10^ to 6.59 x 10^10^ per gram (Fig. 3; Table S5), aligning with viral loads in grassland soils and expected increases post-wet-up (Nicolas et al., 2023). We found that 4.76– 5.15% of total virions were active (Fig. 3; Table S5), representing newly produced particles, while the remainder reflected persistent populations. The persistent virion population was significantly greater than the active virion population across all samples (paired t-test, p < 0.01; Table S5), with a large effect size (Cohen’s d = 3.12), revealing that the majority of the extracellular virions constitute a persistent seed bank rather than newly replicated particles. Mapping stringency had minimal effect on within-sample estimates (Wilcoxon p > 0.05; Table S5), although between-sample differences were significant (Kruskal-Wallis, Dunn’s post hoc, p < 0.05; Table S5), reflecting spatial patchiness and variability in viral DNA recovery typical of soil viromes (Hazard et al., 2025).

**Figure 3.**
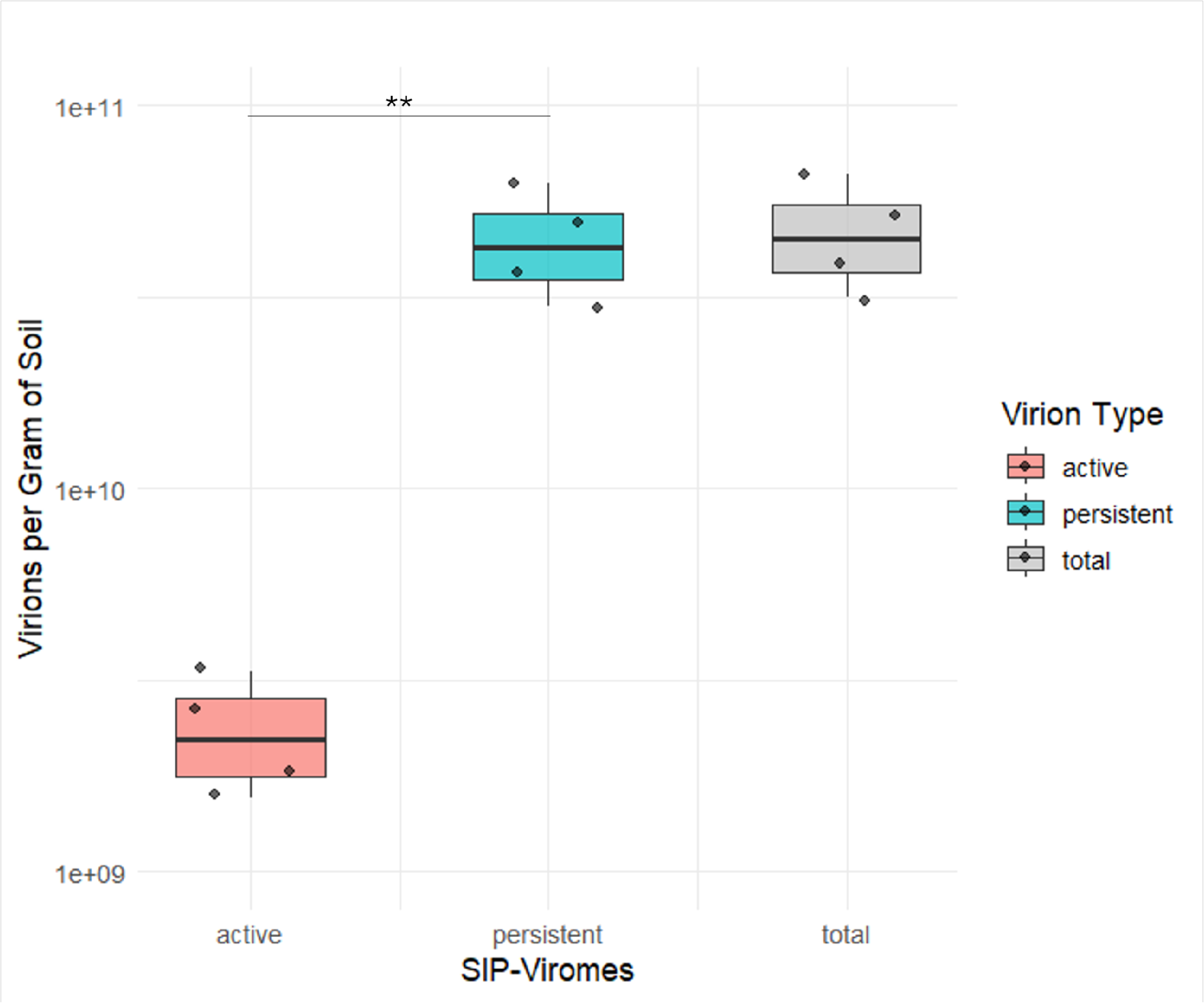
Estimated virions per gram of soil and the proportions that are active or persistent. Three boxplots show the estimated number of active (red), persistent (blue), and the total number of virions per gram of soil for the four soil viromes (10% read mapping threshold). The total number of virions was significantly different across the four soil samples (Kruskal-Wallis test, Dunn’s post hoc, p < 0.05; Table S5), on average, the persistent virion population was significantly greater than the active virion population (paired t-test, p < 0.01; denoted with **, Cohen’s d = 3.12), and estimates across different read mapping stringencies were not significantly different within each sample (Wilcoxon p > 0.05).

To contextualize these empirical estimates and account for the recovery of non-viral DNA (e.g., mobile genetic elements, ultra-small microbes; Trubl et al., 2018, 2020; Nicolas et al., 2021), we performed a deterministic, grid-based sensitivity analysis. This first analysis varied the assumed viral DNA fraction (10–100%) and genome length (minimum, median, maximum from vOTUs), yielding a broad range of potential virion estimates from 4.96 x 10^8^ to 1.03 x 10^11^ per gram of soil (Table S5; Fig. S4). These results represent boundary conditions that help bracket the uncertainty in DNA source attribution and viral genome sizes. Building on this, we conducted a probabilistic Monte Carlo simulation to explore the full distribution of potential outcomes by randomly sampling from a continuous distribution within the same parameter ranges. This approach yielded a range of 1.18 x 10^9^ to 1.01 x 10^11^ virions per gram. Notably, while the full range of the simulated distribution is broad, the central tendency of the simulated distribution closely overlaps with the empirically derived values, indicating that our primary estimates are robust to key sources of uncertainty. This convergence suggests that, although the true virion count may span a broad range due to biological and methodological variability, our experimentally based estimates fall well within the most likely and biologically plausible range defined by the sensitivity and probabilistic analyses.

Host associations and isotopic enrichment

To identify traits associated with virion persistence and activity, we compared ^18^O enrichment across vOTUs with predicted host taxonomy and viral genomic features. Host taxonomy was predicted using iPHoP (Roux et al., 2023), successfully assigning hosts to 32% of vOTUs, spanning five archaeal and twelve bacterial phyla (Table S6). This relatively low rate of host prediction is a common limitation in environmental viromics, especially in soils as there is the highest diversity of viral and microbial genomes and many vOTUs likely represent novel viruses infecting poorly characterized microbial taxa (Roux et al., 2023). Consequently, our interpretation of host-associated patterns is limited to the characterized fraction, and we focus our analysis on the most abundant host predictions, which represent the dominant and well-studied microbial phyla in this system. The most frequent host predictions were Actinomycetota (43%), Pseudomonadota (25%), and Bacteroidota (8%), which are all taxa previously identified as abundant and active in these soils (Greenlon et al., 2022; Nicolas et al., 2023; Foley et al., 2023). Among isotopically enriched vOTUs, 46% (36 vOTUs) had predicted hosts, most commonly Actinomycetota (35%), Pseudomonadota (29%), and Bacteroidota (22%) (Fig. 4). Host phylum significantly explained variation in vOTU EAF (ANOVA, p < 0.001), with a large effect size ∼0.12, although no pairwise differences were significant (Tukey’s post hoc test, P > 0.05), suggesting broad but non-specific host influences on viral persistence. Further analysis showed vOTUs with elevated EAF generally had higher relative abundance (Fig. S5, Spearman ρ = -0.142, p = 0.0058) and were predicted to infect Actinomycetota and Pseudomonadota. Across all vOTUs, EAF showed no consistent relationship with genome length or GC content (Spearman p > 0.05; Fig. S6). These findings indicate that viral replication dynamics in this system are shaped more by host physiology than by viral genomic traits, with the caveat that this conclusion primarily applies to the fraction of the viral community with predicted hosts.

**Figure 4.**
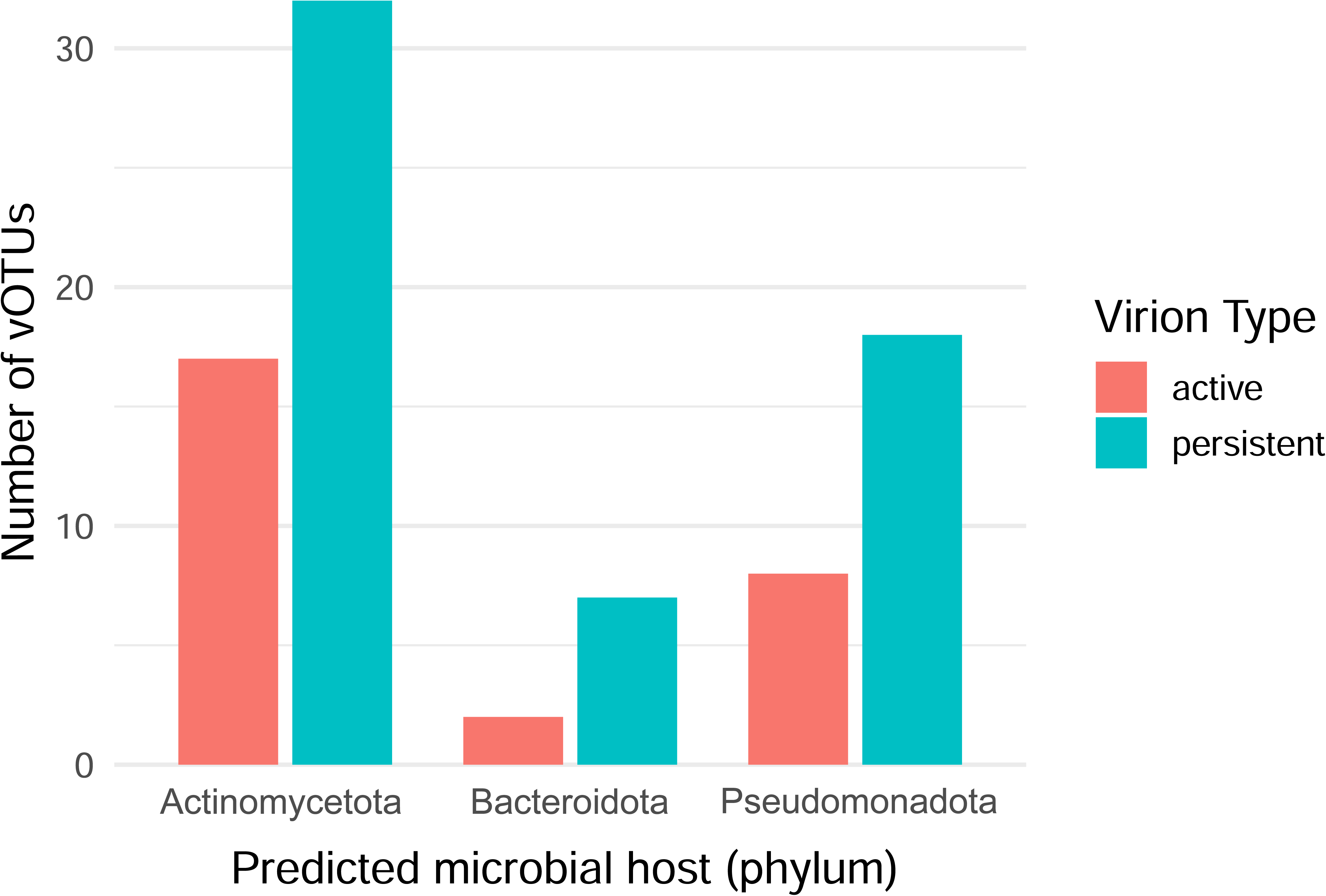
Number of active and persistent vOTUs predicted to infect the top three bacterial phyla. Bar plot showing the number of active (red) or persistent vOTUs (blue) for the top three predicted bacterial hosts at the phylum level.

By integrating ^18^O-SIP with genome-resolved host prediction, SIP-viromics provides a functional, *in situ* readout of lytic infection dynamics. Because extracellular virions can only incorporate ^18^O into DNA following successful replication within a host cell, enrichment of a vOTU whose genome is confidently linked to a host genome is consistent with viral replication occurring within a host from that lineage. Active vOTUs were predominantly associated with Actinomycetota and Pseudomonadota, host lineages that dominate the active microbial community following wet-up in this system, consistent with prior qSIP and metagenomic studies demonstrating that microbial and viral activity co-vary within these lineages following rewetting (Greenlon et al., 2022; Nicolas et al., 2023). This pattern is supported by the isotopic enrichment data (Fig. 2) and by host-resolved counts of active and persistent vOTUs across the top three bacterial phyla (Fig. 4), which show that active vOTUs span multiple host phyla but consistently include Actinomycetota and Pseudomonadota. These observations indicate that viral activity is aligned with host resuscitation dynamics rather than uniformly distributed across all potential hosts. While isotopic enrichment does not distinguish among infection pathways, including replication within low-abundance or unassembled hosts or induction of pre-existing intracellular viral genomes (e.g., proviruses), all of these scenarios require host metabolic activity and result in the production of extracellular virions. Taken together, these findings suggest that viral activity following wet-up is structured by host physiological responses rather than being evenly distributed across the microbial community, highlighting the role of specific host lineages in shaping virion turnover following environmental perturbation. These results further demonstrate that SIP-viromics extends beyond distinguishing newly produced and persistent virions to linking viral activity with host lineages, providing ecological resolution into the host populations contributing to virion turnover following environmental perturbation.

Building on our previous work which showed that dry soil can harbor a diverse but low-biomass reservoir of virions serving as a viral seedbank (Nicolas et al., 2023), our current findings indicate that most virions persisted without replicating throughout incubation, while a distinct subset became active following wet-up. This pattern aligns with the viral seed bank hypothesis, where viruses remain dormant until favorable conditions, such as increased microbial activity following wet-up, trigger infection (Brum et al., 2015; Hazard et al, 2025; Lennon et al., 2021). The majority of active viruses targeted Actinomycetota and Pseudomonadota, bacterial groups known to rapidly respond to rewetting (Nicolas et al., 2023; Santos-Medellín et al., 2023). Notably, nearly 70% of vOTUs predicted to infect Actinomycetota were not isotopically labeled, suggesting their persistence in dry soil prior to rewetting.

A limitation of this study is the absence of a pre-incubation (T0) virome, preventing direct comparison of virion composition before and after wet-up. Although ^18^O-SIP distinguishes newly produced from non-replicating virions, it does not resolve changes in the persistent pool or differentiate long-term persistence from short-term stability. Despite this limitation, the use of ^18^O-SIP uniquely enables inference of viral replication activity independent of initial abundance, providing a complementary approach to temporal sampling.

Taken together, these findings highlight Actinomycetota and their viruses as key drivers of microbial turnover and nutrient cycling in grassland soils, with viral lysis likely releasing nutrients that support broader microbial response after disturbance. In addition to serving as reservoirs of genetic material, persistent virions may play active roles in microbial food webs. Both protists and bacteria have been shown to consume virions, facilitating nutrient and energy recycling (Brown et al., 2020; DeLong et al., 2023; Godon et al., 2021; Martinez-Martinez et al., 2024). Most persistent vOTUs in our study were predicted to infect Actinomycetota and Pseudomonadota, bacterial groups that dominate dry soils, suggesting their associated virions may be particularly adapted to environmental stability. Such stability is consistent with drought-driven changes in soil microenvironments, where reduced water content concentrates solutes and intensifies selection pressures on microbial communities (Shan et al., 2026).

Virion persistence likely stems from co-evolutionary relationships with their hosts, which possess desiccation resistance traits. The associated virions are thus under selection pressure for long-term environmental stability, which may manifest through two major mechanisms. First, persistence is likely linked to drought-adapted traits of the hosts. In these water-limited soils, the host’s protective structures often provide an extrinsic mechanism for virion stability and persistence in the extracellular environment. This includes Actinomycetota’s use of spore formation, thick cell walls, and extracellular polymeric substances (EPS) production (Costa, Raaijmakers, and Kuramae 2018) that shield them from desiccation and osmotic stress. Viruses embedded within these host-derived EPS matrices, and associated with protected host cells, may benefit from a buffered microenvironment that enhances their structural integrity and persistence. Evidence from biofilm systems shows that viral particles can become entrapped within extracellular matrices, where they remain infectious and protected from environmental stressors, effectively functioning as reservoirs that enhance persistence (Bond et al., 2021; Gao et al., 2026). These observations are consistent with the broader role of EPS in retaining water and buffering embedded biological material from desiccation and environmental stress (Flemming and Wingender, 2010). This is further supported by observations that Actinomycetota maintain greater survival in dry conditions (Hestrin et al., 2022; Kristy et al., 2022). For Pseudomonadota, previous work revealed that drought-stressed taxa upregulated carbon and energy conservation pathways, and increased antioxidant defenses (Ghosh et al., 2025; Breitkreuz et al., 2021).

Second, the viral particles themselves may possess intrinsic structural traits associated with long-term environmental stability. Virion stability is likely governed by both intrinsic viral traits, such as capsid geometry, internal pressure, and genome packaging (Cordova et al., 2003; Buenemann and Lenz, 2007; Stone et al., 2019; Podgorski et al., 2025) or specialized proteins that resist shear forces and degradation, promoting virion survival until host dormancy is broken. Capsid architecture, including complexity, directly impacts stability and capacity. For example, decreased icosahedral complexity is predicted to lead to a more stable capsid assembly (Stone et al., 2019). Notably, in Actinobacteriophages, stabilization proteins have been shown to bridge split hexamers in the major capsid proteins, and the length of this stabilization protein adapts to accommodate larger genomes and higher internal pressures, reinforcing the capsid without changing its overall size (Podgorski et al., 2025). The high proportion of persistent vOTUs infecting these drought-adapted hosts suggests a strong selection pressure for virions that are structurally optimized for long-term survival in the soil matrix (e.g., small, highly rigid capsids).

Virion stability is also likely governed by external environmental factors including soil moisture history, temperature, mineral composition, and pH. For example, drying and rewetting cycles can create rapid osmotic pressure changes that lead to the destabilization and disruption of viral capsids (Coleman et al., 2024), and virion decay rates are affected by humidity and droplet size, which are relevant to soil pore environments (French et al., 2023).

Some viruses have evolved structural changes to cope with these stresses, including temperature extremes. These changes appear as unique capsid architectures and modifications to subunit connectivity that enhance capsid stability under environmental stress, such as topological linkages that stabilize the capsid and help it resist high internal and external pressures (Stone et al., 2019). Such structural features are observed across diverse viral lineages and are not restricted to extreme environments (Kim et al., 2019). Furthermore, enhanced hydrophobic interactions at the capsid protein interfaces contribute significantly to the high stability required for survival in harsh environments (Stone et al., 2019). Additionally, virus-mineral interactions, modulated by pH and mineralogy, play a role in virion adsorption (Goyal and Gerba, 1979). Some viruses form occlusion bodies or persist in biofilms, where they remain viable and infectious, both of which can enhance long-term survival under fluctuating soil conditions (Secor et al., 2015; Williams, 2023; Gao et al., 2026). Together, these findings suggest that virion turnover and ecological function in soils are governed by a combination of biotic interactions and abiotic processes that determine environmental longevity.

Our SIP-enabled, host-resolved view of soil viral activity shows that only a small but consequential subset of viruses responds to rewetting, and that these responses are host-specific rather than community-wide. By directly tracing isotope-enriched virions and linking them to predicted hosts, we demonstrate that viral activity following disturbance is constrained by host physiology, local community structure, and recent environmental history, rather than simply by the size of the extracellular viral pool. These results move soil viral ecology beyond presence-absence surveys toward a mechanistic understanding of which viruses’ matter, when, and through which hosts they influence biogeochemical fluxes. In doing so, they complement emerging syntheses that highlight unresolved questions surrounding soil viral abundance, activity, and turnover (Hazard et al., 2025), as well as broader seed-bank frameworks that emphasize temporal storage, delayed feedbacks, and ecological memory across biological systems (Lennon et al., 2021).

The maximum ^18^O enrichment observed in this study was moderate (EAF ≈ 0.25), indicating that the sensitivity of SIP detection was likely conservative and increasing the probability that some actively replicating vOTUs remained below the enrichment threshold. Such underdetection would bias estimates toward overrepresenting persistent virions rather than inflating apparent activity. However, several lines of evidence indicate that this limitation does not alter the main conclusions. The difference between active and persistent virions was large, enriched vOTUs exhibited non-random associations with host taxa known to rapidly resuscitate following wet-up, and virion abundance estimates were stable across deterministic and probabilistic sensitivity analyses. Taken together, these observations suggest that the proportion of active vOTUs reported here represents a conservative lower bound, and that persistent virions dominate the extracellular viral pool following rewetting even when accounting for potential underdetection of viral activity.

Understanding how viruses respond to environmental perturbations, such as soil drying and rewetting, is critical for predicting microbial mortality, nutrient release, and downstream ecosystem processes. Our results show that viral persistence and host-specific replication represent distinct but coupled strategies that shape microbial turnover following disturbance. While persistence reflects the capacity of virions to buffer communities through unfavorable conditions, replication outcomes are strongly shaped by host physiology, viral traits, and abiotic context, consistent with trait-based and assembly-focused perspectives on viral ecology (Brum et al., 2015; Flynn et al., 2021; Liu et al., 2023). Together, these findings indicate that soil carbon and nutrient cycling models must explicitly account for viral infection strategies, host physiological state, and environmental drivers, rather than treating viruses as a uniform mortality term. By identifying the conditions and host lineages under which soil viruses transition from persistence to active turnover, this work establishes a predictive framework linking viral ecology to microbial resilience and ecosystem function under environmental change.

## Materials and Methods

### Sample collection and processing

The soil was collected during the late dry season (August 19, 2019), when field moisture was already low and representative of seasonal drought conditions at an elevation of 1,066 feet (∼325 m) and a depth of 0–10 cm at Buck field site at University of California Hopland Research and Extension Center (GPS coordinates N39°00.106’ W123°04.184’). No additional drying was applied following collection. After sieving using a 2.0 mm wire mesh, 1 kilogram of soil was homogenized and transported back to Lawrence Livermore National Laboratory, where it was stored at room temperature (∼23°C) until being processed the next day. Gravimetric soil water content was measured by drying duplicate 5 g samples to a constant weight at 105°C for 48 hours (Table S5). Soil pH was determined by adding 25 mL dH_2_O to 10 g soil followed by 1 h shaking at 400 rpm and resting for 1 h prior of measuring pH with a sensION+ pH meter (Hach) (FAO, 2021). Four 64-oz (1.89 L) mason jars were filled with 10% HCl overnight to remove organic compounds and triple washed with dH_2_O. Soil was divided up so that 210 g of soil went to each mason jar and then two mason jars received 39 mL of dH_2_O and two jars received 39 mL of heavy water (H_2_^18^O; isotopic purity was 98 atom % ^18^O; ISOFLEX San Francisco, CA, USA).

Water was added to each sample slowly and evenly across the soil surface with a syringe to bring the soil up to 30% moisture (gravimetric water content), mixed, sealed with lids that had airtight septa, and incubated in the dark at room temperature (∼23°C) for 7 days. The samples were harvested on August 26^th^ and soil was divided as follows: 5 grams for pH measurement, 5 grams for gravimetric water content, and 200 grams for a virome. For the virome, the 200 g of soil was divided among 40 50-mL conical tubes with approximately 5 g in each tube and stored at −80°C until processed.

For respiration measurements, 1 mL of headspace was collected from each jar to measure CO_2_ concentration at 0, 1, 3, 5, and 7 days using an air-tight syringe with a 22-gauge needle. The head space was injected into evacuated and sealed Wheaton bottles and stored at room temperature until being measured. CO_2_ from headspace samples was quantified with a LI-850 Gas Analyzer (Licor, Lincoln, NE). Raw data was exported from the Li-850 software and was then analyzed using a custom R script (v4.4.1; https://doi.org/10.5281/zenodo.13875112) to calculate the CO_2_ concentration normalized to soil mass (Table S3; Fig. S2). The data was used to construct a line plot in R using ggplot2 (Wickham 2016) and dplyr (Wickham et al., 2023) and a two-way ANOVA with a Post hoc Tukey test was performed to compare the dH_2_O and H_2_^18^O treatments over the experiment.

Viromes were processed by thawing the tubes at 4 °C for two hours in 10 mL of AKC’ buffer (1% potassium citrate resuspension buffer amended with 10% phosphate buffered-saline, 150 mM magnesium sulfate, 50 µM Na_2_HPO_4_, and 20 µM KH_2_PO_4_; Trubl et al., 2019) and following a previously optimized protocol (Trubl et al. 2016, Trubl et al., 2019) with minor modifications. Briefly, the soil was shaken horizontally for 15 min at 400 RPM followed by 3 cycles of alternating vortexing and hand shaking for 30 s each, then centrifuged at 4 °C at 4,198 x *g* for 20 min. Approximately 50 mL of wash fluid was collected and passed through a Steriflip-GP 0.22 µm filter (Millipore Sigma; cat. # SCGP00525), the resulting filtrate was then concentrated to 250 µL using a 100 kDa Amicon Ultra-15 cellulose membrane filter (Millipore Sigma; cat. # UFC910096) that was pretreated with 3 mL of 1x PBS with 0.05% Tween 20 pH 7.4 (Teknova; cat. # P0201). Additionally, the Amicon filter was washed 3 times with 750 µl of AKC’ buffer per wash. The resulting 2.5 mL of concentrate per sample was then DNase treated with 7.1 µL of RQ1 RNase-Free DNase (Promega, cat. # M6101; Madison, WI) and 270 µL DNase Rxn Buffer (0.1 M Tris-HCL pH 7.5, 25 mM MgCl_2_, 5 mM CaCl_2_) for 30 minutes at 37 °C. Reaction was stopped by adding 135 µL of DNase Stop Buffer (20 mM EGTA pH 8.0; Promega, cat. # M6101; Madison, WI) incubated at 65 °C for 10 min, then stored overnight at 4 °C.

DNA was extracted from virus-like particles following a previously published method (Griffiths et al., 2000), with minor modifications. Briefly, virus-like particles were precipitated using 0.5 mM FeCl_3_ followed by 250 μL of Phenol:Chloroform:Isoamyl (pH 8) added to each sample and vortexed for 1 min. Samples were incubated on ice for 15 min, vortexing occasionally to homogenize samples, and centrifuged at 14,000 g for 5 min. The top layer of the aqueous phase was transferred to a Phase Lock Gel (5 Prime, Gaithersburg, MD, USA) and 250 μL of chloroform was added to the tube. Tubes were centrifuged at 14,000 × g for 5 min, and the supernatant was pooled from each tube and transferred to a fresh tube. Sodium acetate (25 μL, 3 M, pH 5) and 1.5 μL of glycoblue (Invitrogen, 15 mg/mL) were added and mixed and 250 μL of isopropanol was mixed thoroughly with samples and incubated at -80 °C for 20 min. Samples were centrifuged at 14,000 × g for 20 min, and the supernatant was discarded. The pellet was washed with 500 μL of freshly prepared 70% ethanol and centrifuged at 14,000 × g for 5 min. The supernatant was removed from each sample, and samples were centrifuged for an additional 1 min at 14,000 × g. Remaining ethanol was removed from samples and dried for 5 min. DNA was then resuspended in 20 μL 10 mM Tris-HCl pH 8. Resulting DNA concentrate was then quantified (Qubit 2.0; Invitrogen), purity assessed (Nanodrop 200; ThermoFisher) and stored at −80 °C. DNA samples were shipped to JGI for density gradient fractionation, library preparation and Illumina sequencing.

#### DNA density fractionation and sequencing

Before ultracentrifugation and density fractionation, we added a mixture of 9 synthetic DNA oligonucleotides (∼1% of total DNA per sample; Table S7) to each viral DNA extract. These synthetic spike-ins served as fiducial markers along the density gradient to identify potential issues in fractionation (Vyshenska et al., 2023). After adding the spike-ins, the entire DNA sample was loaded into an ultracentrifuge tube and subjected to density gradient centrifugation and fractionation following a previously published protocol (Nuccio et al., 2022) with minor modifications. Briefly, DNA was centrifuged at 44,000 RPM (190,600 × g) for 120 hours in a VTi 65.2 rotor (Beckman Coulter). Forty-eight density fractions of approximately 110 µL were collected from each sample using an Agilent 1260 fraction collector and deposited into 96-well plates. Fraction densities were immediately measured with a Reichert AR2000 refractometer.

To provide an internal standard for normalizing viral coverage levels during later quantitative analysis steps, we added a single synthetic DNA oligo at a fixed mass of ∼100 pg to every density fraction (Table S7). DNA was then pelleted through PEG precipitation, washed in 70% ethanol, and resuspended in 30 µL of TE following Nuccio et al. (2022). Purified DNA concentrations were determined by Qubit fluorometric quantification (Thermofisher). Finally, to reduce the total number of fractions for sequence and increase the amount of DNA per fraction, we combined material from odd-numbered fractions to the adjacent even-numbered fractions (e.g., fraction 1 was added to fraction 2), resulting in 24 total DNA fractions per sample. The density and DNA concentration of the combined fractions was estimated as the average of the two combined fractions.

Fractions with densities of 1.68–1.78 g/mL were selected for sequencing library creation using Nextera XT v2 kits (Illumina) followed by 13 cycles of PCR. Libraries were pooled at equal molar levels based on DNA concentrations measured between 400–800bp on a Fragment Analyzer (Agilent). Pooled libraries were size-selected to 400–800bp with a Pippin Prep 1.5% agarose gel cassette (Sage Science) before sequencing on the Novaseq S4 platform (Illumina) using a 2 × 150 bp paired-end strategy.

Reads were processed with the automated JGI QC pipeline (Clum et al., 2021). Briefly, BBDuk from the BBTools package (https://jgi.doe.gov/data-and-tools/bbtools/) was used to trim reads that contain an adapter sequence or homopolymer stretches of five G’s or more at the ends of reads, quality trim reads where quality drops to zero, and remove reads that contain four or more “N” bases, or have an average quality score across the read of < 3, or have a minimum length of ≤ 51 bp or 33% of the full read length. Reads that can be mapped with BBMap from BBTools to masked human, cat, dog, mouse, and common microbial contaminants at 93% identity are also separated and not included in the assembly process. For each sample, filtered reads of all fractions were then error-corrected with tadpole (mode=correct ecc=t prefilter=2) and deduplicated with clumpify (dedupe subs=0 passes=2) both from the BBTools package (https://jgi.doe.gov/data-and-tools/bbtools/), before assembly using SPAdes v3.13.1 (- -sc --only-assembler -k 21, 33, 55, 77, 99, 127) (Prjibelski et al., 2020), following a pipeline we previously optimized for metagenomes based on PCR-amplified libraries (Roux et al., 2019). The same libraries were also used as input for a combined assembly, performed with MetaHipMer2 version 2.1.0.90 (Hofmeyr et al., 2020).

To further assess non-viral contamination in virome libraries, we applied ViromeQC v1.0 (Zolfo et al., 2019) using raw virome reads as input. Analyses were performed using the environmental preset (-w environmental) with default parameters and reference database, including a minimum read length of 75 bp and a minimum average Phred quality score of 20. Reads were aligned against curated databases of SSU rRNA, LSU rRNA, and single-copy bacterial marker genes using Bowtie2 and DIAMOND as implemented in ViromeQC. ViromeQC is a validated framework that quantifies virome enrichment relative to metagenomes based on the abundance of ribosomal RNA genes and universal bacterial marker genes. Alignment rates were used to calculate an enrichment score relative to unenriched metagenomes, providing a multi-marker proxy for host-derived DNA contamination that extends beyond single-gene (e.g., 16S rRNA gene) estimates. Resulting alignment rates and enrichment scores are reported in Table S2. Each contig set was screened for viruses using VirSorter 2 v2.0 (Guo et al., 2021) using the dsDNAphage, RNA, and ssDNA models, and a minimum score cutoff of 0.5, as well as geNomad v1.5.1 (Camargo, et al., 2023) with a minimum score of 0.7. All predictions from VirSorter 2 and geNomad were then refined through CheckV v0.8.0 (Nayfach et al., 2021), specifically excluding all predicted virus sequences with more than 5 host genes detected, and more than twice as many host than viral genes, based on CheckV quality_summary file. CheckV also provided estimated completeness for each virus sequence prediction. Predicted viral sequences ≥10 kb were then clustered into 354 vOTUs, using the standard cutoff of 95% average nucleotide identity (ANI) over 85% of the shortest sequence (Roux et al., 2019), using CheckV anicalc and aniclust scripts (Nayfach et al., 2021). Filtered reads were mapped to each sample assembly and to the set of vOTUs using BBmap from the BBTools package (https://jgi.doe.gov/data-and-tools/bbtools/) with default parameters. Host prediction was performed with iPHoP v1.3.3 (Roux et al., 2023) using the default host database Aug_2023_pub_rw and a minimum score of 75. Sample information including quality control, assembly, and virus detection can be found in Table S8.

Criteria for determining isotopic enrichment: Per-fraction coverage data for vOTUs and spike-in standards were analyzed with R package SIPmg (Vyshenska et al., 2023), calculating excess atom fraction (EAF) and 95% confidence intervals (CI) from 10,000 bootstrap comparisons between fraction replicates. vOTUs were considered “isotope enriched” with ^18^O if the lower 95% CI EAF bound was greater than 0. vOTU isotope incorporation was visualized with ggplot2 (Wickham 2016). The vOTUs whose DNA showed no significant ^18^O enrichment were defined as persistent virions, representing the collective pool of pre-existing extracellular particles, including those that are viable but did not infect a host and those that are non-viable.

#### Virion calculations and sensitivity analyses

To estimate total virions per gram of soil (V_g_), we adapted a previously published equation (Nicolas et al., 2023), calculating virion abundance for each individual vOTU i using the vOTU’s assembled genome length and relative abundance (Eq. 1; Table S5). Total virions per gram of soil were calculated as the sum of V_g,𝑖_ across all vOTUs.

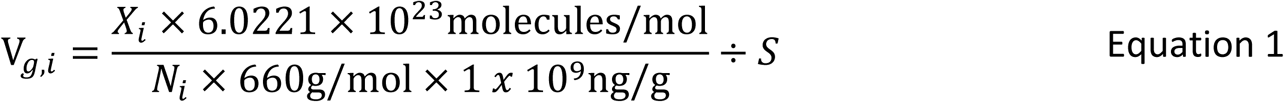

V*g,𝑖*= Virions per gram of soil for vOTU 𝑖

*X*_𝑖_= virome DNA (ng) x normalized relative abundance for each vOTU 𝑖

*N_𝑖_*= genome length of vOTU 𝑖 (bp)

S = grams of soil used for each virion extraction

These calculations assume a relatively constant viral DNA extraction yield across samples and viral genomes/taxa and therefore should be interpreted as order-of-magnitude estimates rather than extraction-yield-corrected absolute abundances. We did not apply an external DNA spike-in correction because free DNA spike-ins do not capture variation in particle recovery. To address these sources of uncertainty, we performed deterministic and probabilistic sensitivity analyses that varied key parameters, including the viral DNA fraction and genome length, thereby bracketing the plausible range of virion abundance estimates rather than assuming a fixed extraction efficiency.

We next calculated the enrichment-adjusted EAF value (EAF_adj_) by normalizing the measured EAF to the atom fraction of ^18^O in the incubation porewater, correcting for dilution of labeled water by native soil water (Eq. 2). This adjustment accounts for underestimation of EAF due to lower than 100 at% tracer available during incubation, providing a more accurate estimate of isotope enrichment.

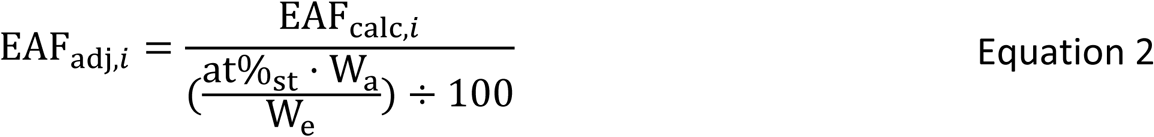

at%_st_ = ^18^O atom percent of the stock water

W_a_ = Volume of water added to each sample (mL)

W_e_ = Total volume of water in each sample during experimental incubation (mL) EAF_calc,𝑖_ = EAF estimates from qSIP calculations

To differentiate between the actively replicating and persistent vOTUs, the total virion abundance is partitioned into two distinct groups (active and persistent). For this study, we are defining ‘persistent virions’ as all pre-existing extracellular particles not undergoing detectable replication during the incubation as indicated by the absence of significant ^18^O enrichment (i.e., EAF 95% CI lower bound ≤ 0) with the acknowledgment that this group could include virions present prior to wet-up that remained viable but did not infect a host, and any non-viable virions that remained detectable as extracellular DNA-containing particles. SIP-viromics cannot distinguish viability, but it can distinguish whether new DNA synthesis and virion production occurred. The abundance of virions that were determined active for each vOTU were adjusted to estimate the new virions produced during the incubation (V_a_, Eq. 3) because the total abundance for each of the vOTUs deemed as active contain both newly produced and persistent virions.

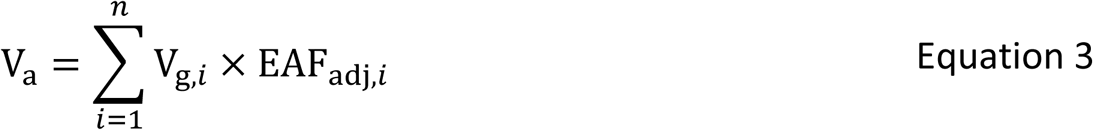

*𝑛* = total number of vOTUs considered in the analysis

To estimate the total abundance of persistent virions (V_p_, Eq. 4), we accounted for two distinct sources of persistence. First, vOTUs classified as non-replicating based on their EAF values were assumed to be entirely persistent, and their total virion abundances were summed directly. Second, for vOTUs classified as active, only a fraction of the population replicated during the incubation; the remaining fraction of these populations was therefore also persistent. The total persistent virion pool was calculated by summing the abundances of vOTUs that never replicated and the non-replicating fraction of vOTUs that were otherwise classified as active, ensuring conservation of total viral abundance. Here, 𝑉_p,𝑖_ and 𝑉_a,𝑖_ represent partitions of 𝑉_𝑔,𝑖_ corresponding to non-replicating and active vOTUs, respectively.

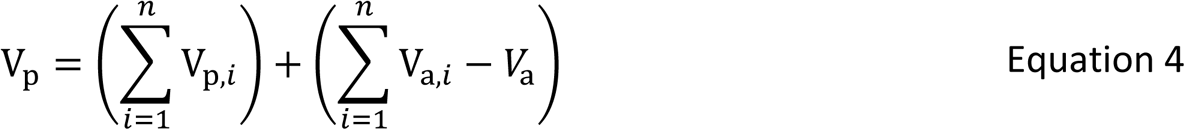

V_p,𝑖_= V_g,𝑖_ for vOTUs classified as non-replicating

V_a,𝑖_= V_g,𝑖_ for vOTUs classified as active

The ‘absolute’ percentage of active virions (V_a_) is determined by calculating the percentage of the total adjusted active virion population compared to the total abundance of virions (Eq. 5). This calculation provides a quantitative metric for the proportion of the total viral community that is actively replicating, serving as a key indicator of viral activity within the soil ecosystem.

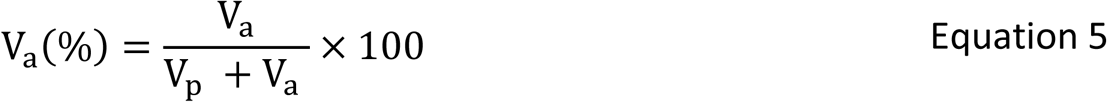

For each sample and mapping threshold (10% and 75% horizontal genome coverage), we computed virions per gram of soil for all vOTUs using their individual genome lengths and relative abundances. We also identified active vOTUs based on isotope enrichment criteria, enabling a comparison of virions derived from active versus persistent populations. To account for biological variability and uncertainty in our assumptions, we conducted sensitivity analyses using both a deterministic and a probabilistic Monte Carlo simulation (Table S5). For the deterministic analysis, we varied two key parameters: the viral DNA fraction and the genome length. For viral DNA fraction, we calculated across a range from 10 to 100% of the total virome DNA originated from viruses, increasing in 10% increments. This approach accounts for the presence of non-viral DNA in viromes, including contributions from mobile genetic elements and ultra-small microbes (Trubl et al., 2018; Trubl et al., 2020; Nicolas et al., 2021). For genome length, we recognized that assembled viral genomes may be incomplete and that true genome sizes can vary, so we estimated virions using the minimum, median, and maximum vOTU genome lengths observed in our dataset. For probabilistic Monte Carlo simulation, we randomly sampled from the same ranges for viral DNA fraction and genome length. Sampling was performed using uniform distributions across these parameter ranges, reflecting equal weighting of all values within the defined bounds in the absence of prior constraints. We performed 1,000 iterations to generate a distribution of potential outcomes, with a random seed set for reproducibility. This approach allowed us to explore the full range of potential virion estimates and determine the most likely virion count distributions, given the uncertainty in these key variables. A R script for calculations of virions, sensitivity analysis, and associated figures is available at: https://doi.org/10.5281/zenodo.16240531.

The applicability and interpretation of Equations 1–5 depend on the isotope tracer used and the degree to which background elemental pools are constrained. Equation 1, which converts DNA mass to virion abundance based on genome length, is independent of isotope chemistry and is transferable across SIP tracers. Equation 2 is also generalizable but must be interpreted as an index of tracer assimilation rather than population growth when applied to ^13^C- or ^15^N-SIP. In contrast, Equations 3–5 rely on the assumption that tracer incorporation reflects total DNA synthesis, an assumption that holds for H_2_ ^18^O but not for ^13^C- or ^15^N in complex soils. Because viruses may replicate using unknown proportions of unlabeled background carbon or nitrogen, these tracers cannot be used to estimate total virion production or persistence. Instead, they quantify the subset of virions produced by hosts assimilating the labeled substrate. Under such conditions, apparent viral ‘persistence’ may reflect tracer decoupling rather than inactivity. Consequently, while ^13^C- and ^15^N-SIP-viromics can provide powerful insights into substrate-specific virus-host linkages, only H ^18^O-SIP enables quantitative partitioning of viral populations into newly produced versus persistent extracellular virions.

---

**Supplementary Figure 1.** Comparison of 10% and 75% coverage profiles of viral populations. (A) Scatter plot showing the relative abundance of each vOTU from stable isotope probing density fractions with the highest concentration (y-axis) plotted against the vOTU’s length (x-axis). (B) Venn diagrams showing the number of distinct or shared vOTUs across the samples based on normalizing the horizontal coverage to 10% or 75% thresholds.

**Supplementary Figure 2.** Soil respiration rates normalized by soil mass over time. The y-axis shows CO_2_ release normalized to soil mass (μg C g⁻¹ soil), and the x-axis shows time (days). Four soil samples were amended with either distilled water or heavy water and monitored over the course of one week. The line represents the sample mean, and shaded area represents the standard deviations (n = 3). Sample CO_2_ measurements are available in Table S3.

**Supplementary Figure 3.** Number of active and persistent vOTUs per sample at both mapping thresholds. Bar plot showing the active (red) or persistent vOTUs (blue) detected in each sample at the 10% and 75% mapping thresholds.

**Supplementary Figure 4.** Sensitivity analysis of virion estimates by viral DNA fraction and genome length assumptions. Estimated virions per gram of soil are shown across a range of assumed viral DNA fractions (10–100%) and three genome length scenarios (minimum, median, and maximum observed vOTU genome lengths). Lines represent individual samples, and colors distinguish samples, while line types group genome length assumptions.

**Supplementary Figure 5.** Relationship between vOTU ^18^O enrichment (EAF) and relative abundance, stratified by predicted host taxonomy. Panels A and B show all vOTUs, with EAF plotted against relative abundance under relaxed (10%, A) and strict (75%, B) enrichment thresholds. Active vOTUs are shown as triangles and persistent vOTUs as circles. vOTUs with predicted hosts are color-coded by host phylum. Panels C and D display only the active vOTUs from A and B, respectively, to highlight trends among active viral populations.

**Supplementary Figure 6.** Pairwise comparisons of vOTU genome features and isotopic enrichment status. Scatterplots, Kernel density plots, and Spearman’s correlation coefficients show relationships among active (red) and persistent (blue) vOTUs, genome length, and GC content.

## Data availability

Data from this project is available at NCBI under BioProject PRJNA1161162, the IMG IDs are included in Supplementary Table 8. The MetaHipMer combined assembly is available on IMG (taxon id: 3300057008).

## Acknowledgments

We acknowledge that the Hopland field site is located on traditional, unceded lands of the Pomo peoples. Research conducted at Lawrence Livermore National Laboratory took place on the territory of xučyun (Huichin), the ancestral and unceded land of the Chochenyo-speaking Ohlone people. We thank Christina Fossum and Rina Estera-Molina for measuring CO_2_ in the headspace of incubation jars, and Tijana Glavina del Rio for coordinating sample shipping and monitoring sample processing at the Joint Genome Institute.

## Funding

This research was supported by a Lawrence Livermore National Laboratory, Laboratory Directed Research & Development grant (18-ERD-041) to S.J.B. and by the U.S. Department of Energy, Office of Biological and Environmental Research (DOE-BER), Genomic Science Program, LLNL “Microbes Persist” Soil Microbiome Scientific Focus Area (#SCW1632). Work conducted at LLNL was conducted under the auspices of the US Department of Energy under Contract DE-AC52-07NA27344. The work (proposal: 10.46936/10.25585/60000473) conducted by the U.S. Department of Energy Joint Genome Institute (https://ror.org/04xm1d337), a DOE Office of Science User Facility, was supported by the U.S. Department of Energy Office of Science, operated under Contract No. DE-AC02-05CH11231.

## Author Contributions

G.T. collected the samples, helped design and conduct the experiment, analyzed the data, prepared the figures, and was the primary writer of the manuscript. S.R. and M.K. analyzed the data, prepared the figures, and contributed to the writing. D.V. performed the DNA density fractionation, analyzed the data, and provided edits on the manuscript. A.T. and K.S. performed the MetaHipMer2 assembly and provided edits on the manuscript. J.A.K. analyzed data and contributed to the writing. E.E-F. helped secure funding for the work and provided edits on the manuscript. R.R.M. helped secure funding for the work, analyzed the data, and provided edits on the manuscript. J.P-R. provided funding for the work, contributed to the writing and provided edits on the manuscript. S.J.B. provided funding for the work, helped design and execute the study, contributed to the writing, provided edits, and served as senior author on the manuscript. All authors helped with the interpretation of the data and approved the final submitted draft.

## Competing interests

The authors declare no competing financial interests.

## Notes

### Competing Interest Statement

The authors have declared no competing interest.

### Summary of Updates

Major revisions were made throughout the manuscript to improve methodological clarity, strengthen interpretation, expand ecological context, and address reviewer feedback. The title was revised from Tracking Persistence and Dynamics of Active Soil Viruses with SIP-Viromics to Disentangling Production and Persistence of Extracellular Virions in Grassland Soils with SIP-Viromics to better reflect major findings and quantitative framework. The Introduction, Results, and Discussion were substantially expanded to clarify the conceptual distinction between newly produced versus persistent extracellular virions, improve interpretation of viral activity following wet-up, and place the work within broader ecological and viral seed-bank frameworks. Additional discussion was added on virion persistence mechanisms, host physiological traits, extracellular polymeric substances, drought adaptation, capsid stability, and environmental controls on virion longevity. The revised manuscript also includes expanded discussion of methodological limitations, including uncertainty in host prediction and interpretation of isotope enrichment pathways. The manuscript now includes extensive methodological revisions and new analyses. These include (i) explicit virion abundance equations and sensitivity analyses, (ii) deterministic and probabilistic Monte Carlo simulations, (iii) clarification of isotope enrichment calculations and assumptions, and (iv) new virome quality-control analyses using ViromeQC to assess non-viral contamination in virome libraries. Additional supplementary figures and tables were added, including ViromeQC outputs, virion sensitivity analyses, active versus persistent virion estimates, and expanded host-association analyses. Interpretation of host associations was revised substantially to clarify that host linkages inferred from iPHoP are probabilistic lineage-level associations rather than direct confirmation of infection of a specific host population. Language throughout the manuscript was updated to reflect this nuance while retaining the central finding that viral activity is strongly associated with dominant, rapidly resuscitating bacterial lineages following wet-up.

