## Supplemental Figures 1-6 for "Disentangling Production and Persistence of Extracellular Virions in Grassland Soils with SIP-Viromics"

Supplementary Figure 1

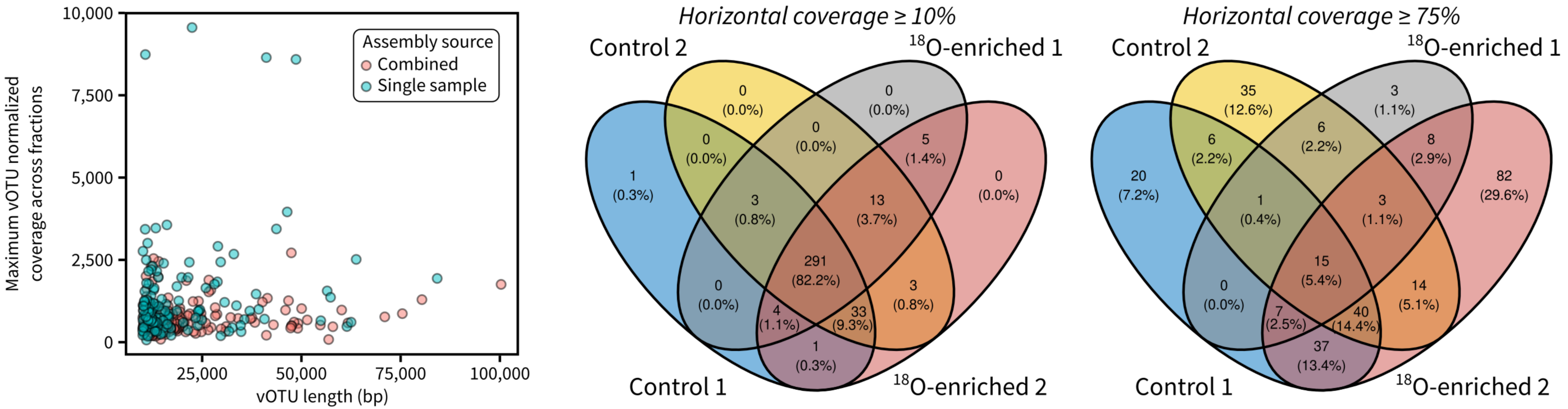

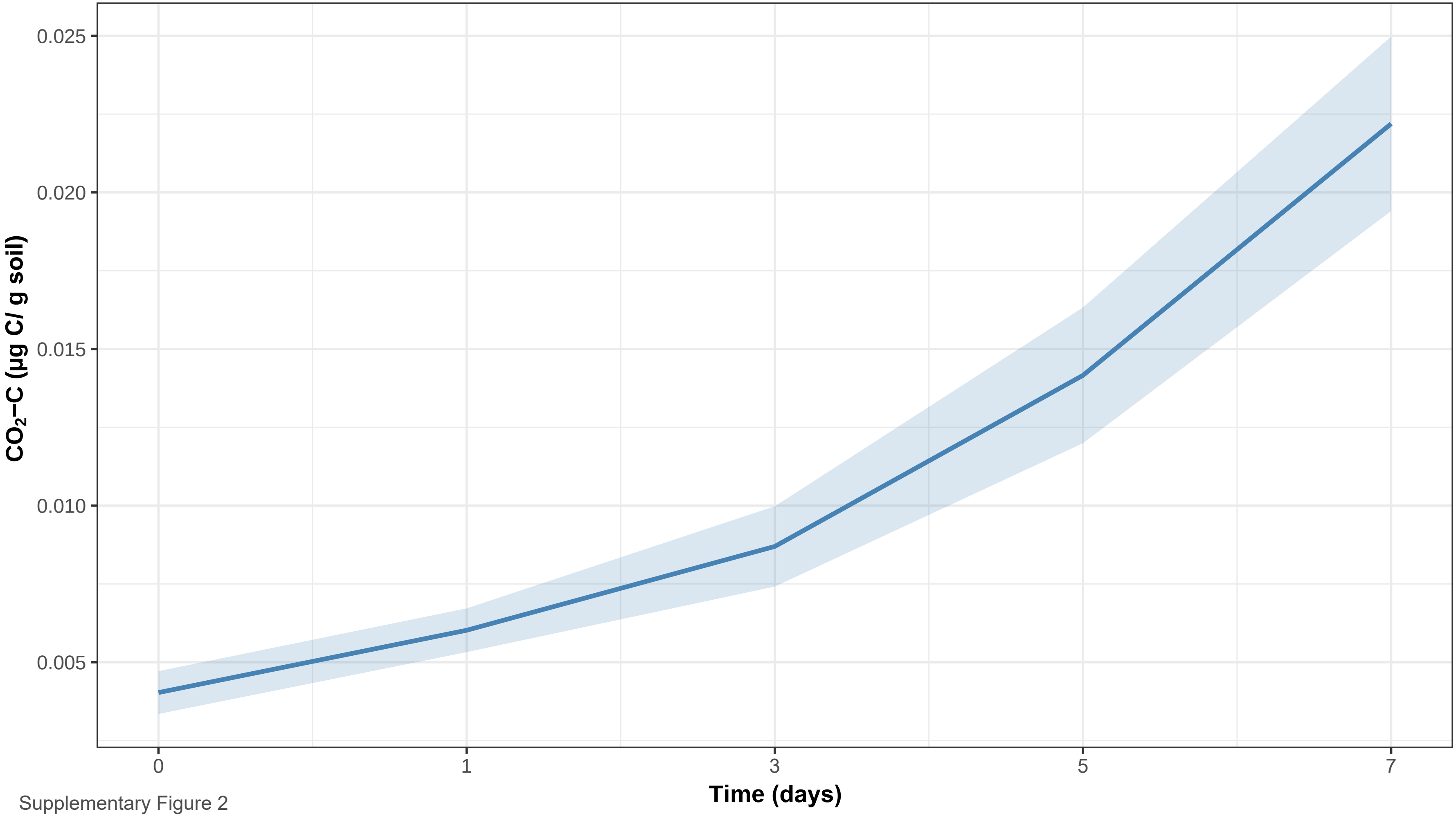


Supplementary Figure 2

Supplementary Figure 3

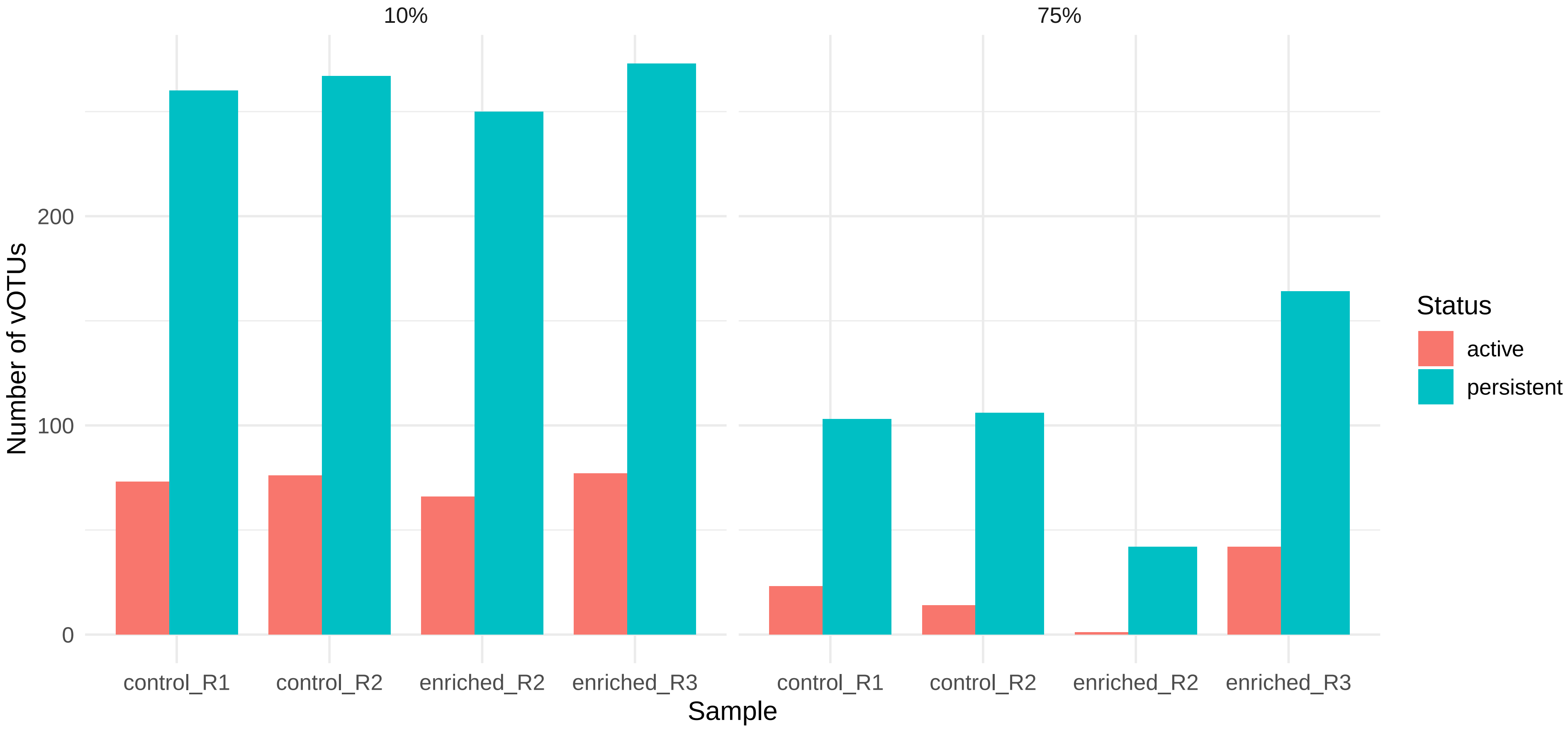

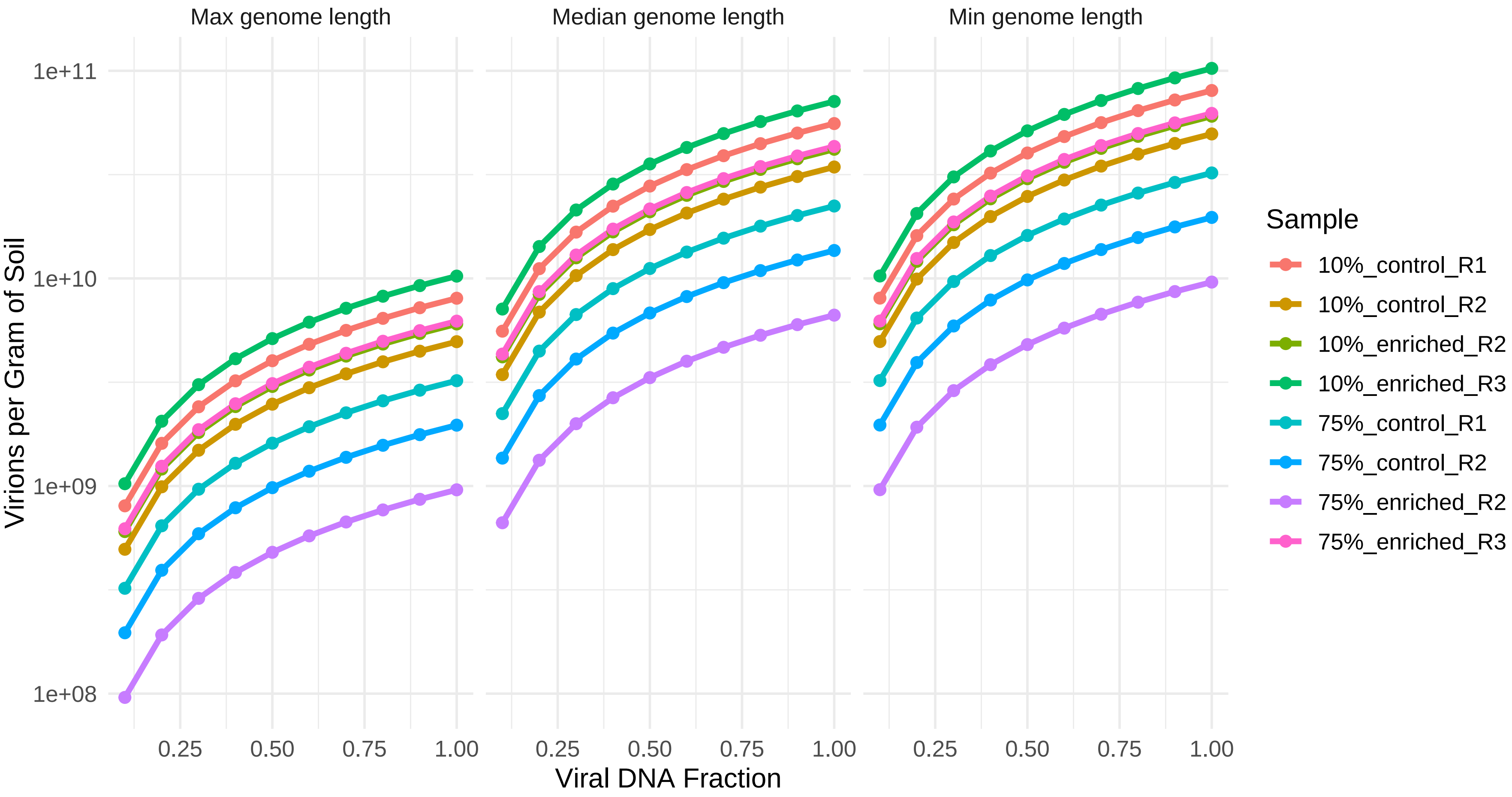


Supplementary Figure 4

Supplementary Figure 5

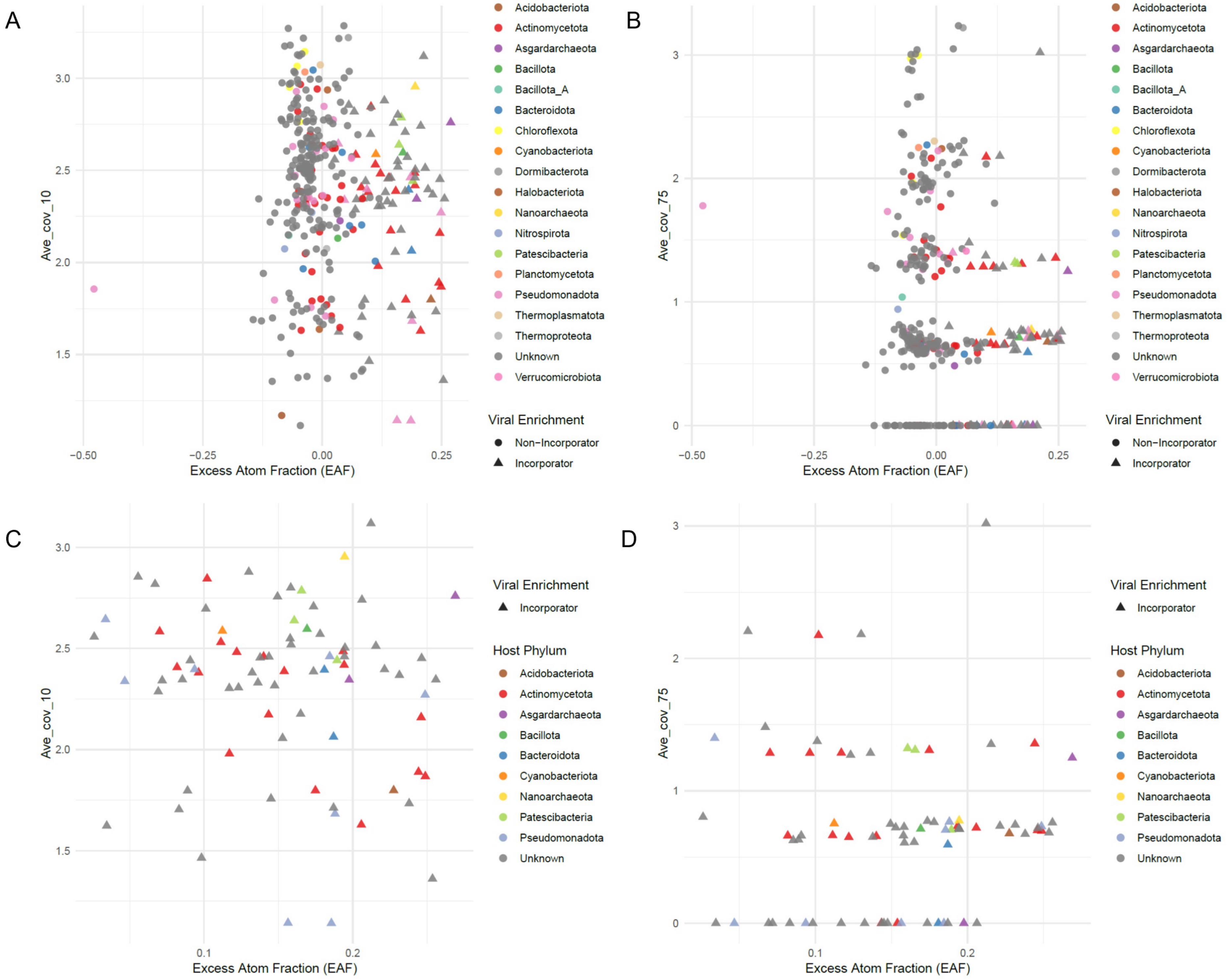

Supplementary Figure 6

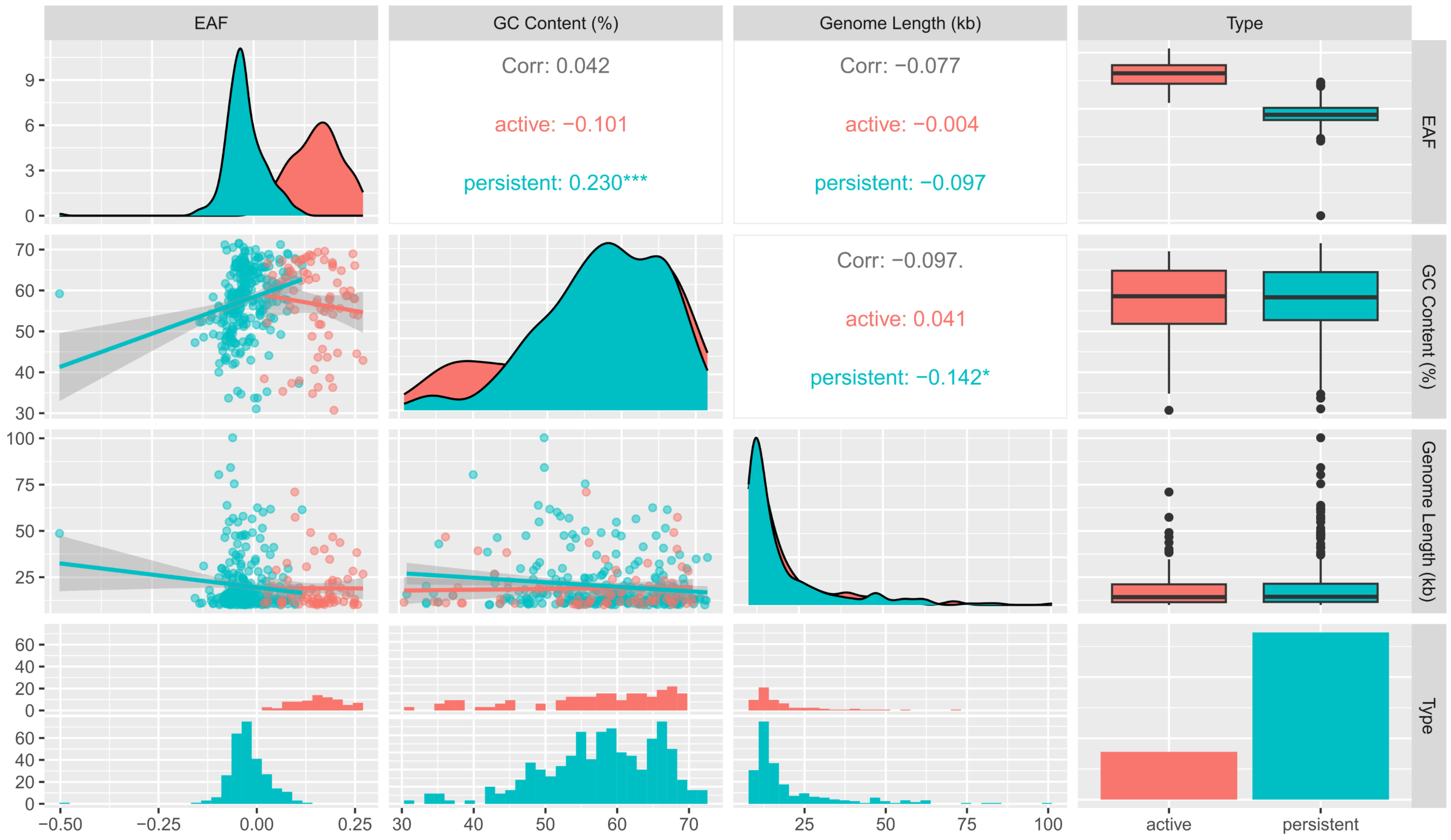
